# Gibberellin and the miRNA156-targeted *SlSBPs* synergistically regulate tomato floral meristem activity and fruit patterning

**DOI:** 10.1101/2023.05.08.539891

**Authors:** Leticia F. Ferigolo, Mateus H. Vicente, Joao P. O. Correa, Carlos H. Barrera-Rojas, Eder M. Silva, Geraldo F.F. Silva, Airton Carvalho, Lazaro E.P. Peres, Guilherme B. Ambrosano, Gabriel R. A. Margarido, Robert Sablowski, Fabio T.S. Nogueira

## Abstract

Many developmental processes associated with fruit development take place at the floral meristem (FM). Age-regulated microRNA156 (miR156) and gibberellins (GA) interact to control flowering time, but their interplay in subsequent stages of reproductive development is poorly understood. Here, we show that GA and miR156 function in tomato FM and fruit patterning. High GA responses or overexpression of miR156 (156OE), which leads to low levels of miR156-targeted *SQUAMOSA PROMOTER BINDING PROTEIN– LIKE* (*SPL/SBP*), resulted in enlarged FMs, defects in FM determinacy and fruits with increased locule number. Conversely, low GA responses reduced fruit indeterminacy and locule number, and overexpression of a miR156-resistant *SlSBP15* allele (*rSBP15*) reduced cell number and size in the FM, as well as locule number. GA responses were partially required for the fruit defects observed in 156OE and *rSBP15* plants. Transcriptome analysis and genetic interactions revealed shared and divergent functions of miR156-targeted *SlSBPs, PROCERA/DELLA* and the classical *WUSCHEL/CLAVATA* pathway, which has been previously associated with meristem size and determinacy. Our findings reveal that the miR156/*SlSBP*/GA regulatory module is deployed differently depending on developmental stage and create novel opportunities to genetically fine-tune aspects of fruit development that have been important for tomato domestication.

## INTRODUCTION

A wide variety of plant organ sizes and shapes exist in nature, and this variation can be explained in part by the control of the apical meristem activity. In maize, for instance, variation in meristem activity is responsible for differences in kernel row number between inbred lines, and high row number is a major driver of yield in cultivated hybrids (Doebley, 2004; Bommert et al., 2013). Regulation of meristem activity occurs at several levels, including identity and determinacy; for example, floral meristem (FM) identity specifies that the meristem can initiate floral organs, and in most cases the FM is determinate, i.e. it is consumed in the production of a limited number of organ primordia (Bartlett and Thompson, 2014). The determinacy and size of the FM set the number of cells available to initiate carpel primordia, and consequently carpel number and ovary size (van der Knaap et al., 2014; Heidstra and Sabatini, 2014). Thus, variation in floral meristem activity in response to hormonal and genetic pathways has had an important impact on fruit patterning and the control of seed dispersal, which have been pivotal to crop domestication and improvement (Purugganan and Fuller, 2009; Østergaard, 2009).

A classic genetic pathway involved in the regulation of meristem size is based on a feedback circuit comprising the stem cell–promoting *WUSCHEL* (*WUS*) homeodomain transcription factor and *CLAVATA* (*CLV*) signaling peptides (Stahl and Simon, 2010). This circuit is conserved in diverse plants, including tomato (*Solanum lycopersium*). Larger meristems in tomato mutants with altered *CLV*-*WUS* activity produce fruits with a higher number of locules (the interior cavities of fruits). Importantly, natural mutations in *CLV3* and *WUS* were essential for tomato domestication (Muños et al., 2011; Xu et al., 2015; Zsögön et al., 2018). Whilst most wild tomatoes and small-fruited cultivars have bilocular fruits, large-fruited cultivars can have eight or more locules (Tanksley, 2004). Most of this variation is due to two mutations (*locule number* or *lc* and *fasciated* or *fas*) with synergistic effects on the number of locules and thus fruit size (Lippman and Tanksley, 2001; Barrero and Tanksley, 2004). The *lc* mutation is localized downstream of tomato *WUSCHEL* (*SlWUS*), whereas a regulatory mutation repressing *SlCLV3* expression underlies the *fas* mutant (Muños et al., 2011; Xu et al., 2015). However, whilst the core *CLV*-*WUS* circuity is deeply conserved, recent work has shown that the pathway is modulated by diverse inputs (Rodriguez-Leal et al., 2019). Given the complexity of genetic and hormonal interactions that regulate meristem function (Lee et al., 2019), there is much potential for discovering novel molecular mechanisms that affect fruit development.

Meristem activity is also regulated by the microRNA156 (miR156) and its transcription factor targets, which are members of the *SQUAMOSA PROMOTER-BINDING PROTEIN LIKE* (*SPL*/*SBP*) gene family. This evolutionarily conserved pathway was initially found to regulate age-dependent processes (Morea et al., 2016). More recently, *Arabidopsis* miR156-targeted *SPLs/SBPs* were shown to regulate the size of both root and shoot meristems (Fouracre and Poethig, 2019; Barrera-Rojas et al., 2020). Down-regulation of *SPLs*/*SBPs* in miR156-overexpressing plants (156OE) also led to abnormal fruit development in both *Arabidopsis* and tomato (Silva et al., 2014; Xing et al., 2013), but with distinct phenotypic consequences: whilst miR156-overexpression only reduced the size of *Arabidopsis* gynoecia, tomato 156OE gynoecia displayed extra carpels and fruit-like ectopic structures (Silva et al., 2014; Xing et al., 2013). Interactions with other pathways also differ between *Arabidopsis*, in which miR156-targeted *SPL* genes control gynoecium patterning through interference with auxin homeostasis and signaling, and tomato, where miR156-targeted *SBPs* modulate the expression of genes involved in meristem maintenance (*LeT6/TKn2*) and organ boundary formation (*GOBLET, GOB*). Thus, similar miRNA-based pathways control initial steps of gynoecium patterning but with distinct functional consequences in dry and fleshy fruit-bearing plants (Correa et al., 2018).

Another molecular player in meristem and fruit patterning is the phytohormone gibberellin (GA). Gibberellin-regulated *DELLA* genes control shoot meristem function, modulating the size of inflorescence and floral meristems (Serrano-Mislata et al., 2017). Furthermore, the GA-deficient *ga1-3* mutant displays delayed growth of all floral organs (Yu et al., 2004). GA controls fruit patterning through the interaction between DELLA and the basic helix–loop–helix (bHLH) proteins INDEHISCENT (IND) and ALCATRAZ (ALC), which specify tissues required for fruit opening. In tomato, on the other hand, scarcely any information is available of how GA modulates FM activity and fruit patterning. The loss of *PROCERA* (tomato *DELLA* gene) function in *procera* (*pro*) mutants or GA_3_ application led to meristic changes in the flower (increased number of all floral organs), and occasional fruits with ectopic fruit-like structures (indeterminate fruits), although the phenotypes are not as strong as in 156OE plants (Carrera et al., 2012; Silva et al., 2014).

In *Arabidopsis*, GA signaling and miR156-targeted *SPLs/SBPs* interact during the floral transition: the GA-regulated DELLA protein REPRESSOR OF GA1-3 (RGA) associates with LEAFY (LFY) and the miR156-targeted SPLs/SBPs proteins to promote *APETALA1* (*AP1*) transcription (Yamaguchi et al., 2014). However, the interactions between the GA and miR156 pathways beyond the floral transition have not been reported. Given the similar phenotypic changes observed in flowers and fruits from tomato 156OE plants and *pro* mutant (Carrera et al., 2012; Silva et al., 2014), we hypothesized that miR156-targeted *SlSBPs* and GA signaling also interact to orchestrate tomato FM activity, early gynoecium patterning and fruit development. Here, we show that the GA and miR156 pathways regulate floral meristem size, and they partially interact at molecular and genetic levels to control fruit determinacy and locule number, without primarily relying on modifications in the classical *SlCLV3*-*SlWUS* circuity, thereby revealing a novel mechanism in the control of tomato fruit patterning.

## RESULTS

### MiR156 and gibberellins have synergistic effects on fruit determinacy

Because we showed that the miR156/*SlSBP* module interplayed with GA to control tomato floral transition (Silva et al., 2019), and that the rice *mir156abcdfghikl* mutant is hyposensitive to GA (Miao et al., 2019), we reasoned that the strong fruit indeterminacy observed in 156OE plants (Silva et al., 2014) could be associated with increased GA signaling. To initially test this conjecture, we quantified bioactive GA_1_ and GA_4_ levels in tomato floral primordia. Whilst GA_1_ levels were undetectable in both WT and 156OE, GA_4_ levels were significantly higher in 156OE primordia compared with WT (Fig. S1A). This finding is consistent with the lower levels of several bioactive GAs (including GA_4_) found in the *mir156abcdfghikl* mutant (Miao et al., 2019). Next, we treated WT and 156OE plants with commercial gibberellic acid (GA_3_). Unlike Carrera et al. (2012), we did not observe indeterminate or malformed fruits in GA_3_-treated WT plants (Fig. S1B). This divergence may be a result of the differences in the GA_3_ treatment (Carrera et al., 2012; Silva et al., 2019). Nonetheless, externally applied GA_3_ led to a 40% increase in indeterminate fruits in 156OE plants compared with mock-treated 156OE plants, suggesting that in addition to higher GA levels, 156OE plants were more sensitive to gibberellin compared with WT (Fig. S1B). Conversely, treatment of 156OE plants with PAC (a GA biosynthesis inhibitor; Jung et al., 2012) resulted in ∼2-fold more normal fruits than Mock-treated 156OE plants, and almost no indeterminate fruits (Fig. S1C).

To evaluate in more detail how gibberellins modulate fruit development in the miR156-overexpressing plants, we inspected ovaries at anthesis and fruits from the hypomorphic *pro* mutant (*pro* harbours a mutation in the *PROCERA/DELLA* gene that lessens protein activity; Carrera et al., 2012; Livne et al., 2015), *GA20ox*-overexpressing plants (GA20oxOE, which show high levels of bioactive GAs; García-Hurtado et al., 2012), and 156OE plants. Neither *pro* nor GA20oxOE plants showed obvious modifications in the ovaries, although 156OE ovaries displayed partially fused extra carpels, as previously described (Fig. 1A-D; Silva et al., 2014). Around 3% of *pro* and 20% of 156OE fruits showed some degree of indeterminacy, as indicated by the development of one or more additional fruit-like structures at the style end of the fruit. On the other hand, GA20oxOE plants did not produce any malformed or indeterminate fruits (Fig. 1I, J). Based on these observations, we hypothesized that high levels/responses of GA can generate strong fruit indeterminacy when *SPLs/SBPs* are lowly expressed in reproductive primordia, as shown in GA_3_-treated 156OE fruits (Fig. S1).

**Fig. 1.**
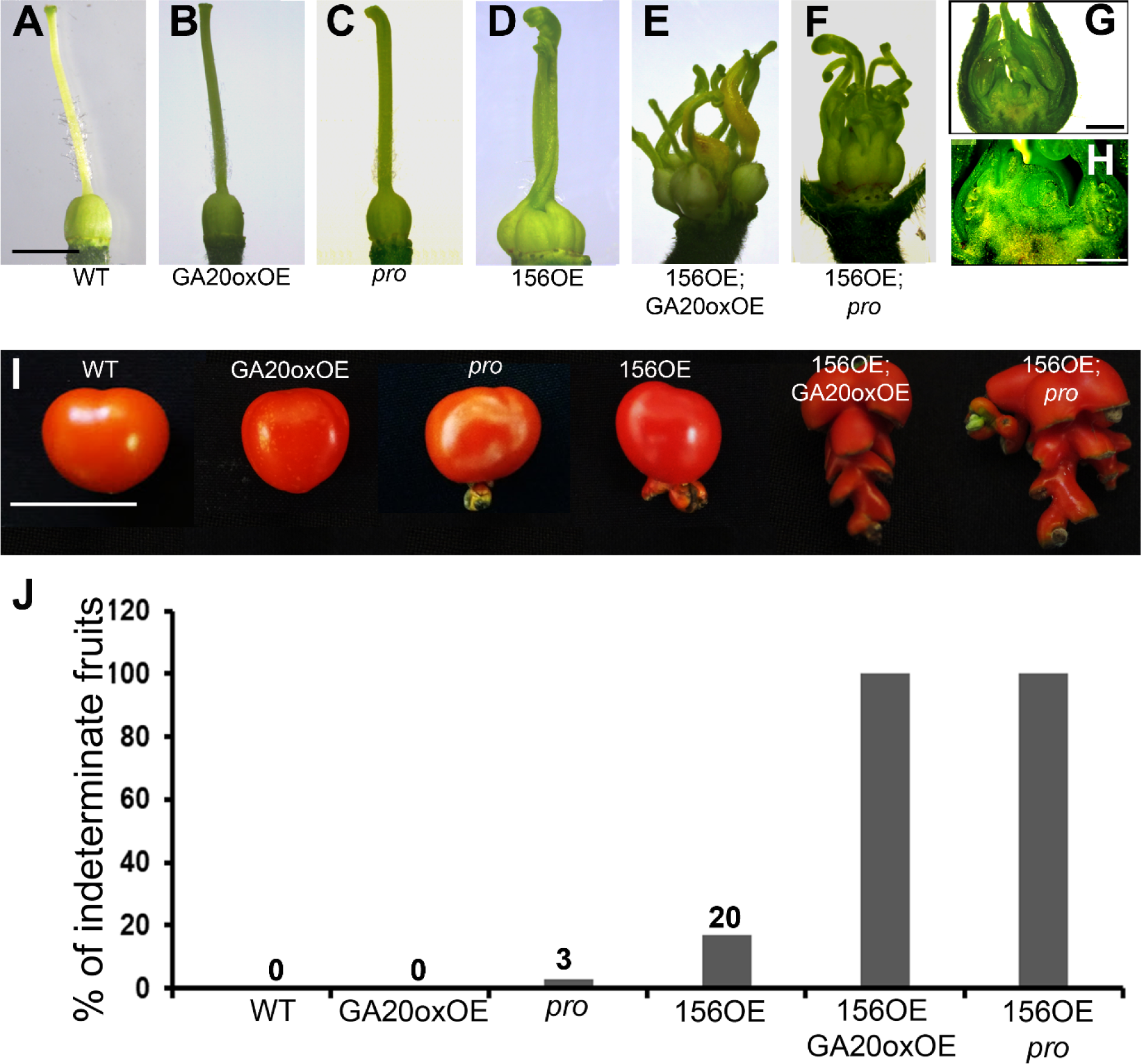
The miR156/*SlSBP* module and gibberellin synergistically regulate tomato gynoecium and fruit patterning. **A-F**) Representative ovaries at anthesis from WT, miR156-overexpressing plants (156OE), *procera* (*pro*) mutant, and plants overexpressing *GA20ox* (GA20oxOE). Scale bar = 2 mm. **G**) Representative flower bud from the 156OE;*pro* plants. Scale bar = 1 mm. **H**) Closeup of the flower bud showed in (**G**). Scale bar = 500 µm. **I**) Representative fruits at 30 days post-anthesis (DPA). Scale bar = 3 cm. (**J**) Percentage of indeterminate fruits (*n* = 120 fruits per genotype).

To genetically confirm the results obtained with GA_3_ treatment, we evaluated ovary growth and fruit patterning in 156OE;GA20oxOE and 156OE;*pro* plants. Strikingly, both 156OE;GA20oxOE and 156OE;*pro* plants showed 100% of fruits with strong indeterminacy (Fig. 1E-J). 156OE;GA20oxOE and 156OE;*pro* flowers display gynoecia formed by supernumerary, partially fused carpels and ectopic pistil-like structures, which did not generated any noticeable locule-like structures (Fig. 1E-H). As a result, the 156OE;GA20oxOE and 156OE;*pro* amorphous fruits were seedless (Fig. 1I). Notably, all these gynoecia and fruits phenotypes were reminiscent of strong miR156-overexpressing tomato lines, which also produced amorphous, seedless fruits (Silva et al., 2014). Collectively, our results indicate that the indeterminate fruit phenotypes observed in 156OE;GA20oxOE and 156OE;*pro* plants are at least in part a result of the synergistic effects of GA and the miR156/*SBP* module converging on the regulation of meristem determinacy and floral organ identity.

### Gibberellins and miR156-targeted *SPLs*/*SBPs* **control locule development and floral meristem size**

Recent evidence indicates that exogenous gibberellin treatment enhances locule number in tomato fruits (Li et al., 2019; 2020). Considering that increased locule number may result from extranumerary carpels due to a mild increase in FM indeterminacy, we checked how the interaction between GA and the miR156/*SlSBP* module affects locule number. In line with the link between indeterminacy and increased locule number, *pro*, GA20oxOE, and 156OE fruits all displayed more locules than WT. Whilst most WT fruits displayed two to three locules, the majority of *pro* and GA20oxOE fruits showed three to five locules (Fig. 2A, B). GA20oxOE and *pro* fruits displayed comparable number of locules, indicating that modifications in either GA levels or responses similarly affect locule development. Consistently, PAC-treated WT plants produced almost 60% of fruits with only two locules and 21% of fruits with three locules (Fig. 2C). The increasing in locule number of *pro* and GA20oxOE was comparable, but not as severe as in 156OE plants, in which most fruits exhibited four to six locules (Fig. 2A, B; Silva et al., 2014). We next inspected locule number in fruits from Mock- and PAC-treated 156OE plants. PAC-treated 156OE plants showed a modest reduction in the number of fruits exhibiting five to seven locules (Fig. 2D), suggesting that miR156-targeted *SlSBPs* and GA control tomato locule number through partially independent mechanisms.

**Fig. 2.**
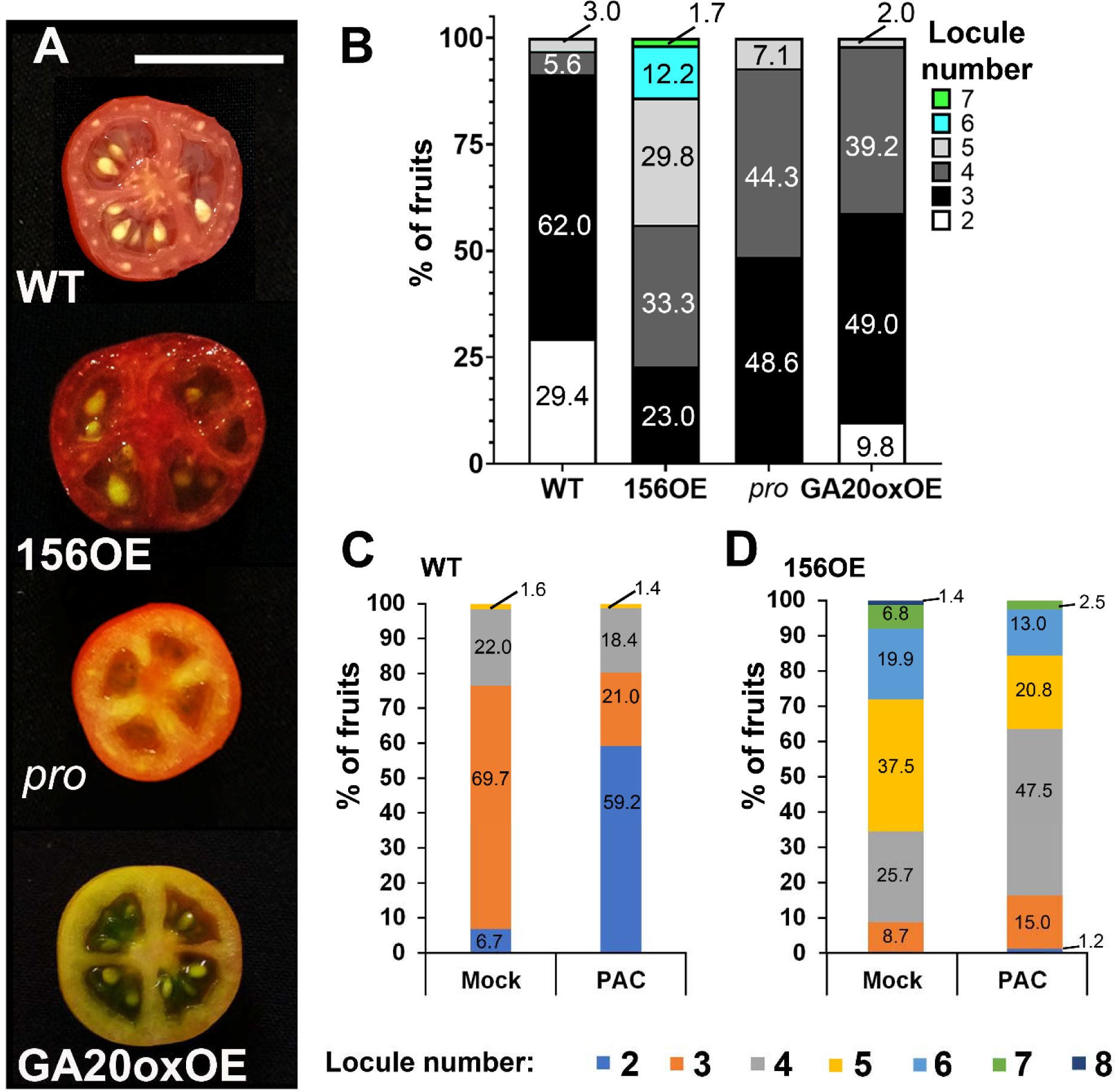
The miR156/*SlSBP* module and gibberellin regulate locule number. **A**) Representative opened 30-day-post anthesis (DPA) fruits from WT, miR156-overexpressing plants (156OE), *procera* (*pro*) mutant, and plants overexpressing *GA20ox* (GA20oxOE). Scale bar = 2 cm. **B**) Percentage of fruits producing distinct number of locules in each genotype. (*n* =150 fruits/genotype). **C, D**) Percentage of fruits exhibiting distinct number of locules in Mock (ethanol)- and 10^-6^ M of paclobutrazol (PAC)-treated WT (**C**) and 156OE (**D**) plants. (*n* = 100 fruits/genotype).

Meristem determinacy and size are tightly correlated with the rate of organ initiation, as larger meristems produce more organs per unit time, including carpels (Xu et al., 2015; Je et al., 2016; Serrano-Mislata et al., 2017). Because *pro* and 156OE plants displayed variable degrees of fruit indeterminacy and locule number (Fig. 1, 2), we compared floral meristem size in WT, *pro* and 156OE plants. To precisely measure FM size, we first established in tomato the modified pseudo-Schiff propidium iodide (mpSPI) and imaging methodology described for *Arabidopsis* inflorescence meristems (Bencivenga et al., 2016; Serrano-Mislata et al., 2017) (Fig. S2). To compare this parameter among different genotypes, we harvested tomato floral primordia with FMs displaying emergence of sepal primordia in a helical pattern (At 1-2 dpi; Xiao et al., 2009). After segmenting and identifying FM cells in 3D, we used the combined volume of cells in the L1 layer of the meristem dome as a proxy for meristem size (Fig. 3A). 156OE exhibited slightly larger meristems than *pro*, and both FMs were significantly larger than WT FMs (Fig. 3B). Importantly, these results correlate FM volume and the number of locules among these genotypes (Fig. 2, 3). To identify the cellular basis for the differences in meristem size, we analysed the cell number and cell volume in the L1 of tomato floral meristems. The mean number of cells in 156OE FMs was significantly greater than WT and *pro*. The cell number of *pro* meristems was intermediate between that of WT and 156OE meristems (Fig. 3C). In addition, the enlargement in floral meristems of 156OE and *pro* (Fig. 3B) was associated with increase cell volumes, although 156OE and *pro* were comparable (Fig. 3D). Our findings indicated that reduced *DELLA/PRO* and *SlSBP* activities enlarged tomato FM by increasing both cell size and number.

**Fig. 3.**
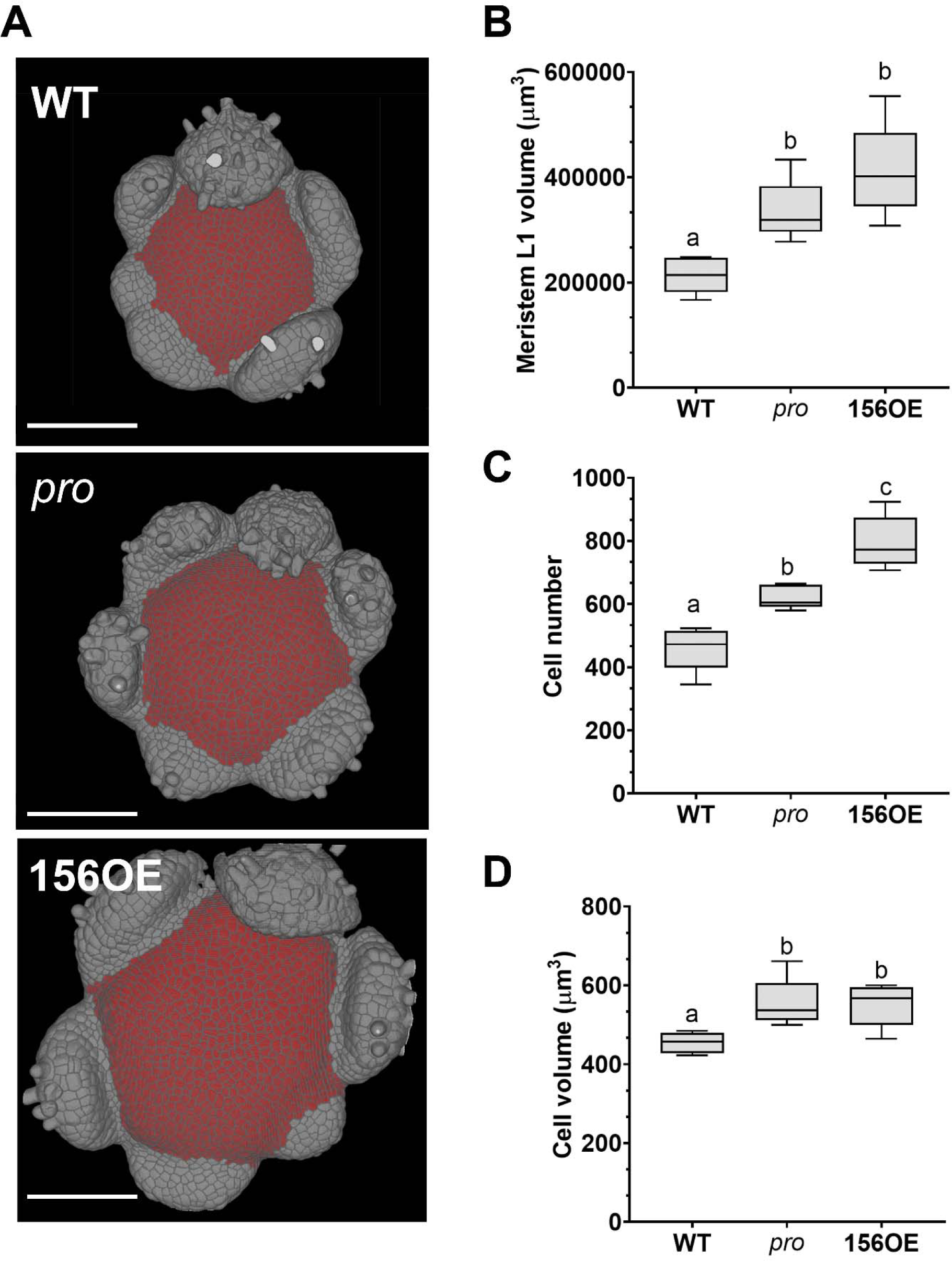
Low activity of miR156-targeted *SlSBPs* and *DELLA/PROCERA* leads to an increase in cell size and cell number in tomato floral meristem. **A**) Top views of 3-D reconstructions of processed confocal stacks of representative floral meristems (FM) from WT, miR156-overexpressing plants (156OE), and *procera* (*pro*) mutant. The FMs were stained with modified pseudo-Schiff propidium iodide (mpS-PI), and the cells highlighted in red were selected as meristem L1 cells. Scale bars = 100 µm. **B-D**) Box plot representations of meristem L1 volume (**B**), cell number (**C**), and cell volume (**D**) (*n* = 5) determined as previously described (Serrano-Mislata et al., 2017). Black center lines show the median; box limits indicate the 25th and 75th percentiles; whiskers extend to 5th and 95th percentiles. Letters show significant differences between genotypes (*P* < 0.05, using ANOVA followed by Tukey’s pairwise multiple comparisons).

### Transcriptional reprogramming of 156OE floral primordia

We have previously shown that some *SlSBPs* and genes associated with boundary establishment (i.e, *GOBLET*) were mis-expressed in 156OE developing ovaries (Silva et al., 2014). To better understand the molecular mechanisms by which the miR156-targeted *SlSBPs* interact with GA responses and how they modulate meristem activity and fruit patterning, we used RNA-seq to monitor changes in gene expression in the 156OE floral primordia. RNA-seq experiments were conducted at 1-2 dpi (which comprise inflorescence meristems and FMs displaying emergence of sepal primordia; Xiao et al., 2009). At these developmental stages in WT plants, miR156 mature transcripts were weakly localized in the flanks of floral meristems, and they were barely detected in the inflorescence meristem (Fig. S3C). In contrast, mature miR156 transcripts were readily detected in the dome of early and late vegetative meristems, as well as in leaf primordia (Fig. S3A, B), consistent with the miR156 role as a master regulator of the vegetative phase (Hyun et al., 2016).

The RNA-seq analysis detected 240 differentially expressed genes (DEGs) between 156OE and WT floral primordia (Table S1). As expected, the DEGs included miR156-targeted *SlSBPs*, which were down-regulated in 156OE primordia. Two strongly repressed *SlSBPs* (3 to 4-fold) were *SlSBP3* (Solyc10g009080) and *SlSBP15* (Solyc10g078700), which are representatives of the two main SPL/SBP clades (Table S1; Fig. 4A; Morea et al., 2016). *SlSBP3* and *SlSBP15* are expressed during *S. pimpinellifolium* early floral development, but with distinct expression profiles (Fig. 4B; http://ted.bti.cornell.edu/cgi-bin/TFGD/digital/home.cgi; Wang et al., 2019). Based on the fruit tissue specific expression atlas (https://tea.solgenomics.net/), *SlSBP3* and *SlSBP15* are preferentially expressed in the pericarp and septum of 0-DPA (days post-anthesis) tomato fruits (Fig. 4C), which may suggest that these *SlSBPs* have roles in orchestrating the patterning of these tissues.

**Fig. 4.**
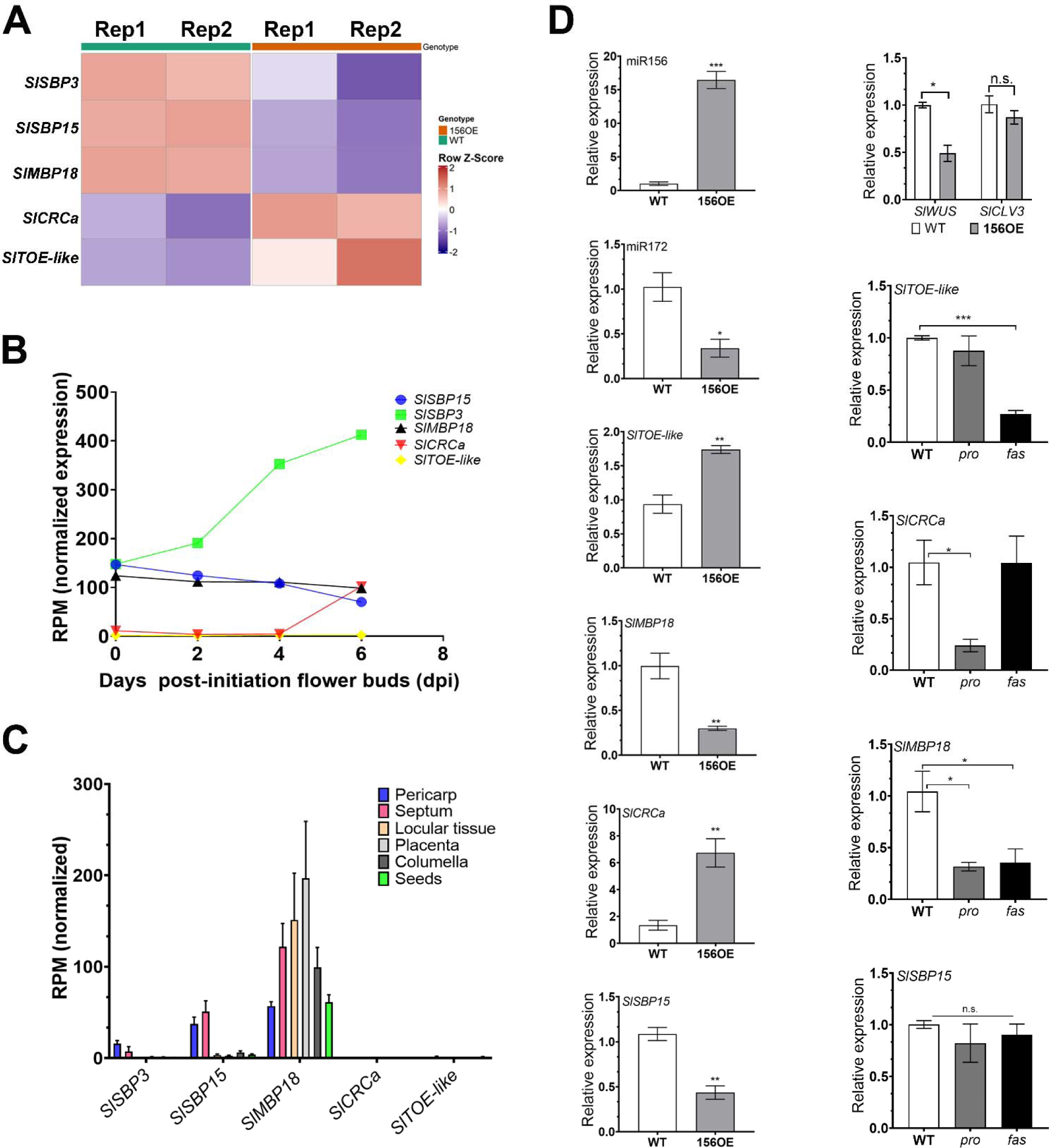
Distinct genetic reprogramming of miR156-overexpressing (156OE) floral primordia. **A**) Heatmap showing the expression profiles (from two replicates of the RNA-seq data) of tomato *TARGET OF EAT-like* (*SlTOE*ℒ*like), CRABS CLAWa* (*SlCRCa*), *SQUAMOSA PROMOTER BINDING PROTEIN–LIKE3* and *-15* (*SlSBP3* and *SlSBP15*), and *MIKC^c^-Type MADS-Box SlMBP18*. **B**) Normalized expression (reads per million, RPM) data retrieved from http://ted.bti.cornell.edu/cgi-bin/TFGD/digital/home.cgi (Wang et al., 2019). **C**) Normalized expression level data (reads per million, RPM) retrieved from the tomato expression atlas (https://tea.solgenomics.net/). Values are mean ± SE. **D**) Relative expression (by qRT-PCR in independent samples) of *WUSCHEL* (*SlWUS*), *CLAVATA3* (*SlCLV3*), *SlTOE*ℒ*like, SlCRCa, SlMBP18* and *SlSBP15* in 1-2 day-post-inflorescence (dpi) WT, 156OE, *procera* (*pro*) and *fasciated* (*fas*) primordia. Values are mean ± SE (n = 3). \**P*<0.05, \*\**P*<0.01, \*\*\**P*<0.001, according to Student’s t-test (two-tailed).

We identified genes directly associated with floral determinacy, such as the tomato *CRABS CLAWa* (*SlCRCa*; Solyc01g010240). Recently, it has been shown that *SlCRCa*, and its paralog *SlCRCb*, are positive, redundant regulators of tomato FM determinacy (Castañeda et al., 2022), so it was surprising that *SlCRCa* was up-regulated in 156OE FM, which has decreased determinacy (Table S1; Fig. 4A, D). However, it has been reported that in the *slcrcb* null mutant, *SlCRCa* transcripts accumulated at higher levels than WT, likely due to a compensatory mechanism activated upon partial loss of determinacy (Castañeda et al., 2022). We speculate that a similar mechanism activates *SlCRCa* in 156OE floral primordia, as 156OE fruits also show a partial loss of determinacy, which can be greatly enhanced, for example in the *pro* mutant background (Fig. 1; Silva et al., 2014).

Another interesting DEG was Solyc04g049800, a *TARGET OF EAT-like* (*TOE*ℒ*like) AP2-type* gene, which is faintly expressed during early flower development and in fruit tissues (Fig. 4B, C) and was up-regulated in 156OE primordia (Table S1; Fig. 4A, D). Solyc04g049800 (*SlTOE-like*) is a target of miR172 (Karlova et al., 2013), which is repressed by the combined action of AtSPL9 and DELLA proteins to regulate the floral transition in *Arabidopsis* (Yu et al., 2012). Moreover, miR172 positively regulates tomato floral identity in a doseDdependent manner via *AP2-like* target genes, including *SlTOE-like* (Lin et al., 2021). This raised the possibility that miR172/*AP2* module might also take part in downstream responses to the miR156/*SlSBPs* in the tomato FM, and indeed, miR172 was down-regulated in 156OE floral primordia (Fig. 4D).

In spite of the strong genetic interaction between 156OE and GA signalling, our RNA-seq data did not reveal changes in the expression of genes involved in GA biosynthesis and signalling associated with reduced expression of miR156-targeted *SlSBPs* (Table S1). This result is consistent with the idea that most integration of miR156-targeted SlSBPs and GA-targeted DELLA proteins occurs by direct protein-protein interactions to regulate shared downstream targets (Yamaguchi et al., 2014). A candidate shared target was the *MIKC^c^-Type MADS-Box SlMBP18* (Solyc03g006830; Hileman et al., 2006), which was down-regulated in 156OE primordia (Table S1; Fig. 4A, D). SlMBP18 is 49% identical to *Arabidopsis AGAMOUS-like 42* (*AGL42*), which promotes *Arabidopsis* flowering in a gibberellin-dependent manner (Dorca-Fornell et al., 2011). In line with a role downstream of miR156-targeted *SlSBPs* in floral primordia, *SlMBP18* was expressed during early flower development in most tissues of 0-DPA fruits, partially overlapping with the *SlSBP3* and *SlSBP15* expression patterns (Fig. 4B, C).

Links to the WUS/CLV pathway were also not apparent in the RNA-seq data: *SlWUS* (Solyc02g083950) and *SlCLV3* (Solyc11g071380) were expressed in our floral primordia samples but were not detected as differentially expressed (adjusted p-values = 0.63 and 0.41, respectively). Neither were other genes associated with the *CLV*-*WUS* pathway such as *WUSCHEL-RELATED HOMEOBOX5* (Solyc06g076000; adjusted p-value = 0.53; Rodriguez-Leal et al., 2019) and *CLAVATA3/ESR (CLE)* genes (Solyc05g009915 and Solyc11g066120, adjusted p-values = 0.84 and 0.80, respectively; Rodriguez-Leal et al., 2019). Considering the central role of *WUS* and *CLV3* in meristem size (Stahl and Simon, 2010), we included *SlWUS* and *SlCLV3* among the genes whose expression in 156OE and WT floral primordia was independently verified by qRT-PCR (Fig. 4D). *SlCLV3* expression did not change significantly, but qRT-PCR did show down-regulation of *SlWUS* in 156OE floral primordia (Fig. 4D), perhaps because the qRT-PCR experiment had higher statistical power. Another possible explanation for this incongruity is that the *SlWUS-SlCLV3* circuity operates at later stages in tomato flower development (Chu et al., 2019; Rodriguez-Leal et al., 2019). However, the lack of changes in *SlCLV3* expression and other genes associated with the *CLV*-*WUS* pathway makes it unlikely that *SlSBPs* targeted by miR156 affect FM meristem size and determinacy primarily through the WUS/CLV pathway.

The results above suggested that the miR156/*SlSBP* module does not regulate FM function by modulating the expression of genes involved in GA signalling or in the *WUS/CLV* pathway. To test whether GA signalling or the *WUS/CLV3* pathway modulate the expression of miR156-regulated *SlSBPs* or their downstream targets, we compared the expression of selected genes in floral primordia of WT, *pro* and in the *fas* mutant, which has reduced *SlCLV3* expression (Barrero and Tanksley, 2004; Chu et al., 2019). MiR172-targeted *SlTOE-like* was down-regulated in *fas* floral primordia, but it was similarly expressed in *pro* and WT. Conversely, *SlCRCa* was expressed at comparable levels in WT and *fas* primordia, but it was repressed in *pro* (Fig. 4D). Neither *pro* nor *fas* affected *SlSBP15* expression, but both mutants had lower expression of *SlMBP18*, as seen with 156OE (Fig. 4D). Overall, our transcriptomic data suggest that the miR156/*SlSBP* module, *PRO* and *FAS* do not regulate each other’s expression, but share overlapping sets of downstream targets during early floral development.

### Activation of miR156-targeted *SlSBP15* reduces meristem activity and attenuates GA effects

Overexpression of miR156 is expected to inhibit *SPL/SBP* expression, but the role of specific miR156-targeted *SlSBPs* in tomato meristem activity and fruit patterning is unclear. Given that *SlSBP3* and *SlSBP15* represent two distinct *SlSBP* clades (Morea et al., 2016) and were strongly down-regulated in 156OE floral primordia (Table S1; Fig. 4), we next investigated how their de-repression would affect tomato FM activity and fruit patterning.

We initially generated tomato Micro-Tom (MT) lines overexpressing a miR156-resistant *SlSBP3* allele (namely *rSBP3*; Fig. S4A). We evaluated three *rSBP3* lines, all of which showed much higher *SlSBP3* transcript levels than WT, and no indeterminate fruits (Fig. 5A, S4B). *rSBP3* lines displayed higher percentages (69 to 95%) of bilocular fruits when compared with WT (Fig. S4C). Given that *rSBP3* lines #2, #4 and #6 displayed similar vegetative architecture and fruit phenotypes (Fig. S4C, D), we selected *rSBP3#2* for further analyses. To study the effects of *SlSBP15* activation, we used a line with overexpression of a miR156-resistant *SlSBP15* (*rSBP15*), which leads to axillary bud arrest and reduced lateral branching (Barrera-Rojas et al., 2022). Like *rSBP3#2, rSBP15* plants produced no indeterminate fruits, and most of the fruits showed an apparent decrease in locule number (Fig. 5A). In summary, the effects of either *SlSBP3* or *SlSBP15* overexpression on fruit development were opposite to those seen in 156OE plants.

**Fig. 5.**
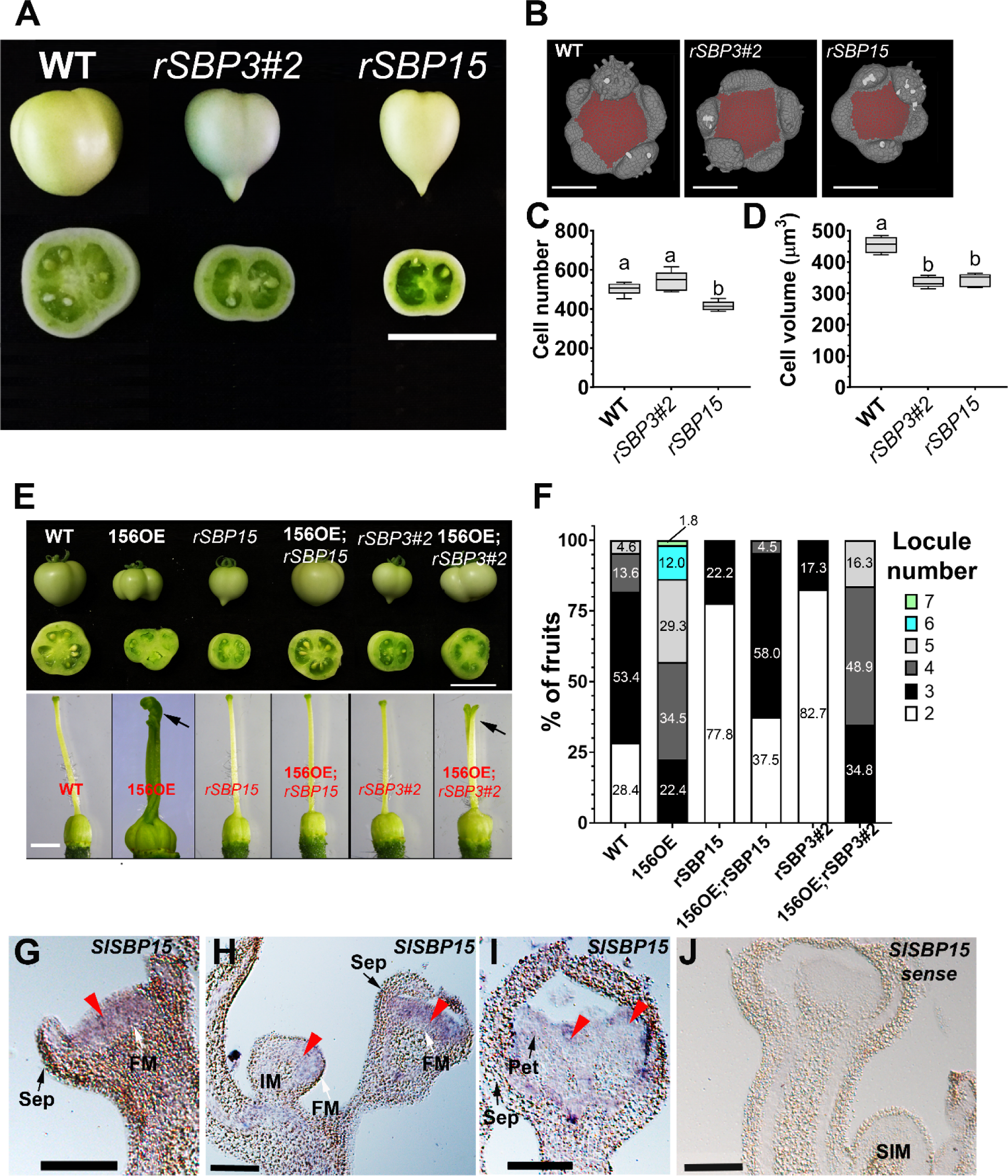
Expression of a miR156-resistant version of *SlSBP15* leads to a decrease in cell size and cell number in the floral meristem, and reduced locule number in fruits. **A)** Representative fruits of WT and plants overexpressing miR156-resistant versions of *SlSBP3* (*rSBP3#2*) and *SlSBP15* (*rSBP15*). Bar = 2 cm. **B**) 3-D reconstruction of processed confocal stacks of representative floral meristems (FM) from WT, *rSBP3#2* (*rSBP3*), and *rSBP15* plants observed from the top. The FMs were stained with modified pseudo-Schiff propidium iodide (mpS-PI), and the cells highlighted in red were considered for quantifications. Scale bars = 100 µm. **C, D**) Box plot representations of cell number (**C**) and cell volume (**D**) (*n* = 5) in FMs determined as previously described (Serrano-Mislata et al., 2017). Black center lines show the median; box limits indicate the 25th and 75th percentiles; whiskers extend to 5th and 95th percentiles. Letters show significant differences between genotypes (*P* < 0.05, using ANOVA followed by Tukey’s pairwise multiple comparisons). Image and data for WT are the same as those in Fig. 4. **E**) Representative fruits (top) and ovaries at anthesis (bottow) of WT, miR156-overexpressing plants (156OE), *rSBP3#2, rSBP15* plants and the double transgenic 156OE;*rSBP3#2* and 156OE;*rSBP15* plants. Bar = 2 cm (fruits) and 2 mm (ovaries). Arrows indicate stigma. Image for 156OE ovary is the same as that in Fig. 2. **F**) Percentage of fruits producing distinct number of locules in each genotype. (*n* =150 fruits/genotype). **G-I**) A digoxigenin-labeled antisense probe detecting *SlSBP15* transcripts was hybridized with longitudinal sections of young inflorescences showing the floral meristem (FM) (**G**), FM plus inflorescence meristem (IM) (**H**), and floral bud at four to five days post inflorescence (dpi), with petals (Pet) emerging over the sepals (Sep) (**I**). Purple staining shows probe localization (red arrowheads). Scale bars: 100 µm. **J**) A digoxigenin-labeled sense probe was used as a negative control. SIM, sympodial meristem.

We next measured *rSBP3#2* and *rSBP15* FM size as described in Fig. 4. The activation of *SlSBP3* only marginally reduced floral meristem size, whereas *rSBP15* meristems were significantly smaller than *rSBP3#2* and WT plants (Fig. 5B, S5A). Cell number in *rSBP15* FMs decreased compared with *rSBP3#2* and WT, which did not differ from one another (Fig. 5C). On the other hand, both *rSBP3#2* and *rSBP15* FMs showed smaller cell volumes compared with WT (Fig. 5D). These results suggest that the functions of different *SlSBPs* targeted by miR156 do not fully overlap and that *SlSBP15* has a more prominent role in FM size.

To evaluate how each of the two *SlSPBs* could restore the functions inhibited by the miR156 overexpression, we crossed the *rSBP3#2* and *rSBP15* transgenes into the 156OE background. Only 34.8% of the 156OE;*rSBP3#2* fruits had three locules, with the majority of the fruits having four or more locules (Fig. 5E, F). Moreover, most 156OE;*rSBP3#2* ovaries at anthesis exhibited partially fused styles, similar to 156OE ovaries (arrows at Fig. 5E). These observations indicate that the activation of *SlSBP3* in 156OE is not sufficient to rescue reproductive defects of 156OE plants. On the other hand, 156OE;*rSBP15* plants showed no indeterminate fruits, and WT-like ovaries at anthesis. Importantly, over 95% of 156OE;*rSBP15* fruits had 2-3 locules, comparable with WT fruits (∼82%) (Fig. 5E, F). Thus, activation of *SlSBP15* restored most of the developmental processes that are disrupted by miRNA156 overexpression. To test whether loss of *SlSBP15* function is sufficient to explain the defects seen in 156OE plants, we examined (CRISPR)-ASSOCIATED NUCLEASE 9 (Cas9) gene-edited *SlSBP15* plants (*sbp15^CRISPR^*; Barrera-Rojas et al., 2022). This loss of function mutant showed no fruit indeterminacy, and most fruits (over 80%) resembled WT (Fig. S5B). Therefore, additional miR156-targeted *SlSBPs* operating in the FM (Table S1) likely function redundantly with *SlSBP15* to control FM activity and fruit patterning.

Because *rSBP15* overexpression led to a stronger reduction in FM size and was sufficient to partially rescue WT-like ovary and fruit phenotypes (Fig. 5E, F), we monitored by *in situ* hybridization the *SlSBP15* expression pattern in early flower development. At 1-2 dpi, *SlSBP15* was mostly detected in flat floral meristems, which showed the emergence of sepal primordia at distinct developmental stages (Fig. 5G, H). By contrast, *SlSBP15* transcripts were scarcely detected in sepal primordia or inflorescence meristems (Fig. 5H). At 4-5 dpi, *SlSBP15* was lowly expressed in floral buds (Fig. 5I). No signal was observed with the *SlSBP15* sense probe (Fig. 5J). These findings reinforced that *SlSBP15* has an important role in controlling FM activity.

*rSBP15* plants exhibited semi-dwarfism (Barrera-Rojas et al., 2022; Fig. S6), a characteristic GA-deficient or GA-insensitive phenotype. Thus, we reasoned that high levels of *SlSBP15* might attenuate GA responses or decrease GA sensitivity in tomato. To test this conjecture, we crossed the *rSBP15* transgene into GA20oxOE plants, which have increased GA levels. In contrast to the typically tall GA20oxOE plants, *rSBP15*;GA20oxOE plants were semi-dwarf like *rSBP15* plants (Fig. S6A). Importantly, *rSPL15* also partially reverted the effects of GA20oxOE in the fruit: *rSBP15*;GA20oxOE plants produced a high percentage of WT-like fruits with two to three locules, in contrast with the 3-4 locules seen in most GA20oxOE fruits (Fig. 6A, B). Thus, *SlSBP15* activation reduced the effects of high GA levels on tomato development. To test to what extent *rSBP15* mimics the effects of reduced GA signalling, we also crossed the *rSBP15* transgene into *pro* background. The resulting *rSBP15*;*pro* plants showed similar vegetative architecture as *pro* plants (Fig. S6B) but had fewer indeterminate fruits than *pro* (Fig. 6C), even though the frequencies of locule numbers were similar (Fig. 6B). In conclusion, the genetic interactions suggest that *rSPL15* opposes GA signalling during vegetative growth and in determining locule numbers but can also compensate for the effects of loss of GA signalling on fruit determinacy.

**Fig. 6.**
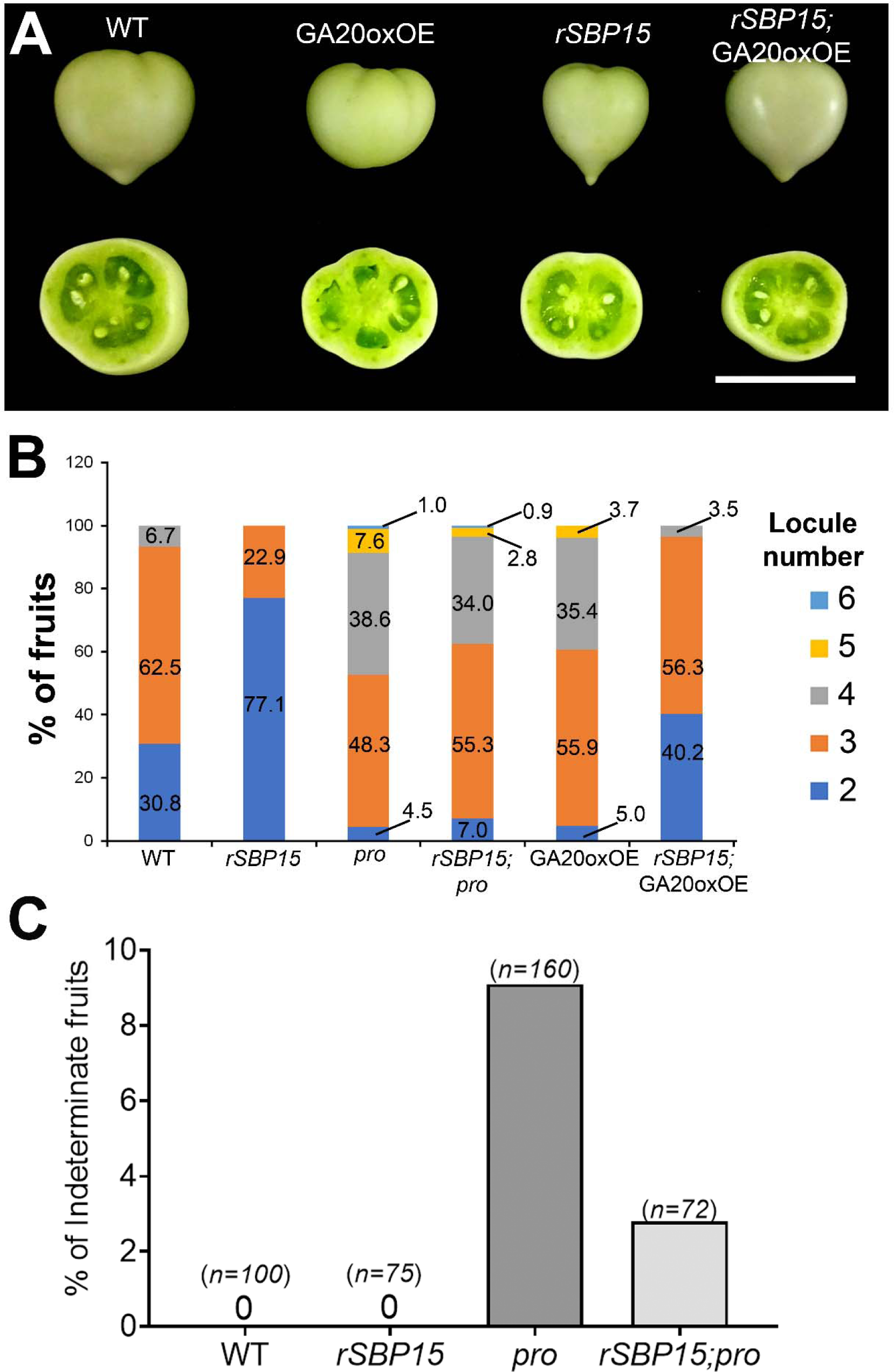
Expression of a miR156-resistant version of *SlSBP15* attenuates gibberellin responses in fruit development. **A**) Representative fruits of WT, miR156-resistant *SlSBP15-*overexpressing plants (*rSBP15*), GA20oxOE, and *rSBP15;*GA20oxOE. showing the locules in each genotype. Scale bar = 2 cm. **B**) Percentage of fruits producing distinct number of locules from WT, *rSBP15, procera* (*pro*), GA20oxOE, *rSBP15;pro* and *rSBP15;*GA20oxOE. (*n* =100 fruits/genotype). **C**) Percentage of indeterminate fruits in WT, *rSBP15, pro*, and *rSBP15;pro* plants. *n* represents the number of fruits/genotype.

### Distinct genetic interactions among the miR156/*SlSBP15* node, *DELLA/PRO* and *SlCLV3* during tomato fruit development

The data presented above suggested that the *miR156*/*SlSBP* module interacts differently with GA signalling during vegetative development, fruit patterning and determinacy. Furthermore, our RNA-seq data suggested that the effects of miR156/*SlSBP* in early FM development are not associated with clear changes in *WUS/CLV3* expression. Like *fas* mutant (Chu et al., 2019), *pro* and 156OE plants also showed enlarged SAM, while *rSBP15* displayed smaller SAM area compared with WT (Fig. S7A, B). We hypothesized that the miR156/*SlSBP* module and *DELLA*/*PROCERA* act along with the *SlWUS-SlCLV* circuity at least partially via common pathways. To investigate how the *WUS/CLV3* pathway interacts with the miR156/*SlSBP* module and GA signalling at the genetic level during fruit development, we next crossed 156OE, *rSBP15* and *pro* with the *fas* mutant.

The combination of 156OE with *fas* showed dramatically enhanced defects in flower development and fruit patterning. 156OE*;fas* plants had supernumerary, partially fused floral whorls such as sepals, petals, and carpels (Fig. S8A) and produced 100% of indeterminate fruits, in contrast to no indeterminate WT fruits, and ∼20% of indeterminate fruits in both 156OE and *fas* plants (Fig. 1J, 7A-D). Amorphous 156OE*;fas* fruits had ectopic fruit-like structures growing from their stylar end, with no apparent locular area (Fig. 7D, S8A), similar to fruits from 156OE;GA20oxOE and 156OE;*pro* plants (Fig. 2).

**Fig. 7.**
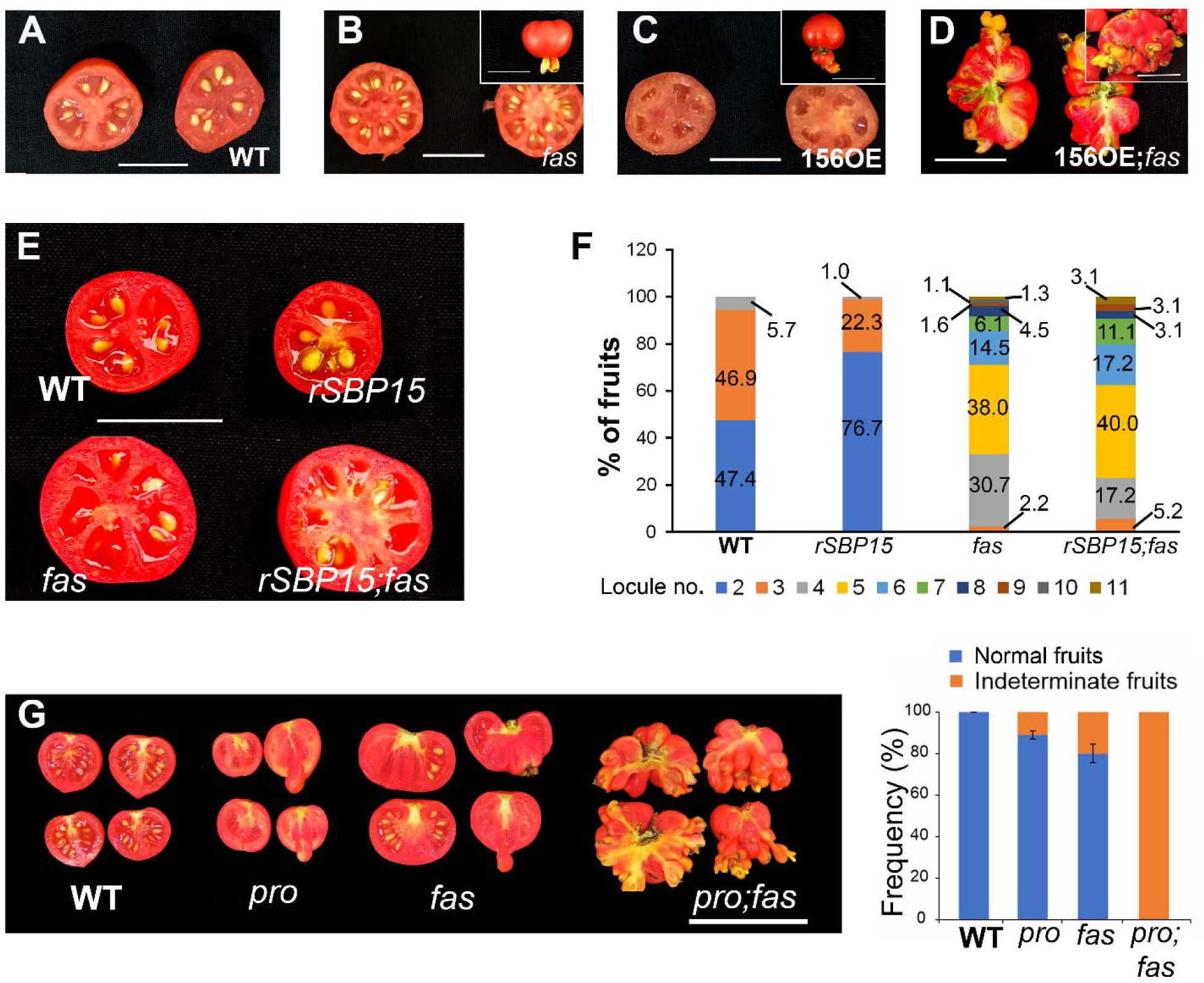
*DELLA/PROCERA* and the miR156/*SlSBP* module act synergistically with *SlCLAVATA3* (*SlCLV3*) to control tomato fruit patterning. **A-D**) Representative fruits at 30 days post-anthesis (DPA) from WT, miR156-overexpressing plants (156OE), *fas*, and 156OE;*fas*. Insets show indeterminate fruits displaying fruit-like structures growing from their stylar end. Bars = 3 cm. **E**) Representative fruits at 30 DPA. **F**) Percentage of fruits producing distinct number of locules from WT, miR156-resistant *SlSBP15-* overexpressing plants (*rSBP15*) and *SlCLV3* mutant *fasciated* (*fas*). (*n* =100 fruits/genotype). **G**) Left panel: Representative fruits at 30 DPA. Bar = 5 cm. Right panel: Percentage of normal and indeterminate fruits. (*n* =60 fruits/genotype).

Conversely, *rSBP15* completely suppressed the fruit indeterminacy seen in the *fas* background (Fig. S7C). Thus, combined loss of *FAS* and *SlSBP* function had synergistic effects on fruit determinacy, and *SlSBP15* activation could bypass the determinacy defects seen in *fas*. However, the genetic interactions differed in relation to locule number: as described before, *fas* mutants showed a large increase in locule numbers, which remained similar in *rSBP15*;*fas* plants (Fig. 7E, F). Thus, as seen above for the interaction with GA signaling, the miR156/*SlSBP15* node and *FAS* interact differently in the control of locule number and fruit determinacy.

We next checked the interaction between *fas* and GA signalling. *Arabidopsis* DELLAs have been reported to restrict inflorescence meristem size independently of the canonical *CLV*-*WUS* circuity (Serrano-Mislata et al., 2017). Similarly, application of the bioactive GA_4_ or the GA inhibitor PAC modified locule number without relying on changes in the expression levels of *SlWUS* or *SlCLV3* (Li et al., 2020), and mutation of the tomato *DELLA* led to enlarged SAMs in the *pro* mutant (Fig. S7A, B). As seen for 156OE, combined loss of *pro* and *fas* had synergistic effects. The flowers from the double *pro;fas* mutant displayed excessive partially fused carpels containing extra carpels within, which were rarely observed in the *pro* or *fas* single mutants (Fig. S8B). While ∼ 9% and 20% of indeterminate fruits were observed in *pro* and *fas* mutants, respectively, *pro;fas* plants produced 100% of indeterminate fruits, all characterized by fruit-like structures growing from their stylar end, and no visible locular area (Fig. 7G, S8B). PAC treatment marginally reduced locule number in the *fas* mutant (Fig. S7D), suggesting that the increased locule number in *fas* is at least partially dependent on GA signaling.

Collectively, our genetic and molecular observations support the idea that miR156-targeted *SlSBPs, DELLA/PROCERA*, and *SlCLV3* have overlapping functions in FM and fruit development, but their interactions vary between different aspects of fruit development, such as determinacy and the regulation of locule number. All the genetic interactions are summarized in the Table 1.

**Table 1.**
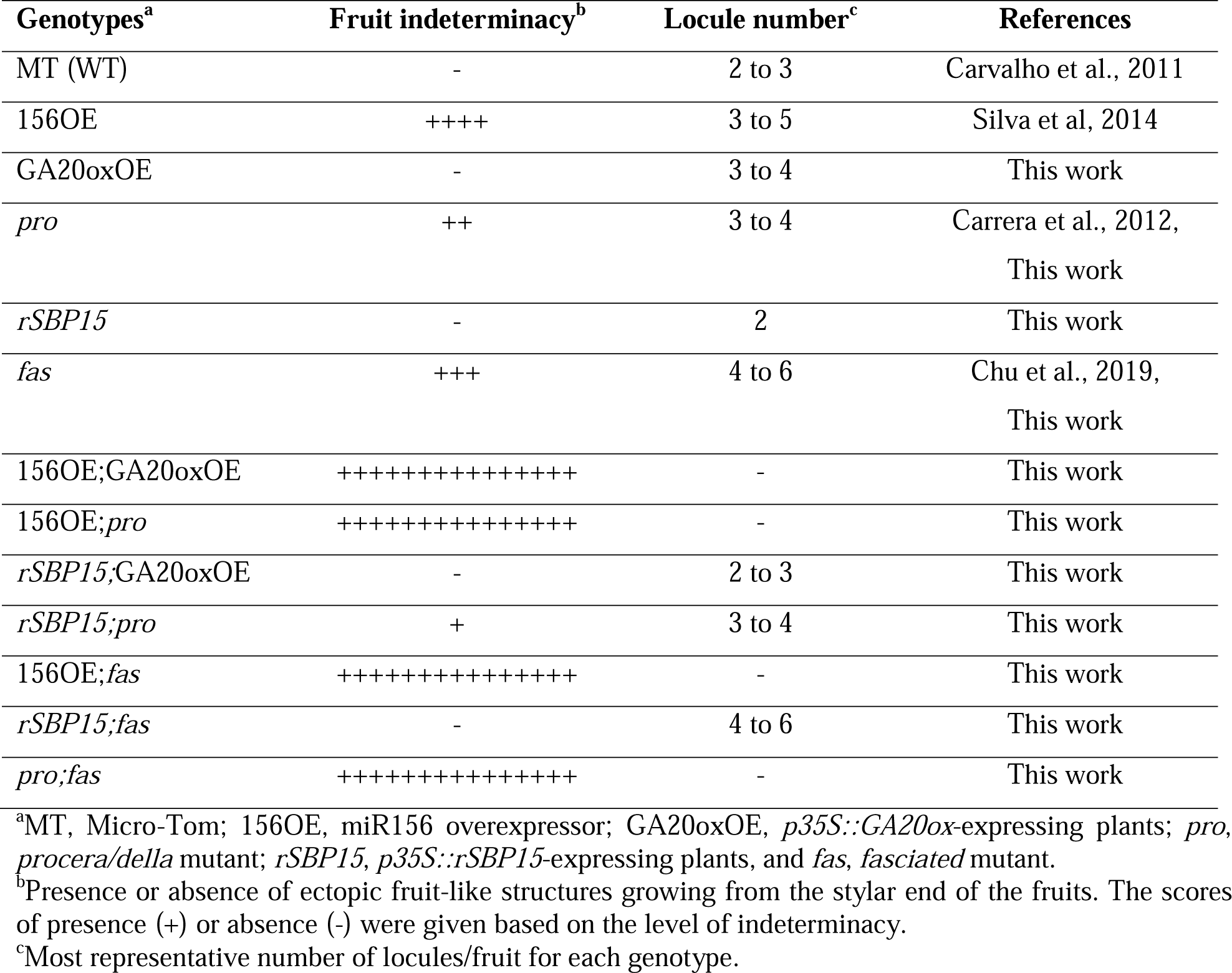
Summarized fruit phenotypes of the main genotypes used in this work.

### *GOBLET* functions later in tomato gynoecium patterning, and it is regulated by the miR156/*SlSBP15* node and *PROCERA/DELLA*

The data presented so far suggested that changes in locule number and fruit determinacy must be under distinct regulation and do not simply unfold from early differences in FM size. One process that may change locule number without necessarily affecting determinacy is the formation of organ boundaries, in which genes of the *CUC* (*CUP-SHAPED COTYLEDONS*) family play a major role. Multiple lines of evidence implicate *GOB* (Solyc07g062840, a homologue of the *Arabidopsis CUC2*) in the control of carpel and locule number in tomato. First, overexpression of miR164, which inhibits *GOB*, reduced locule numbers (Silva et al., 2014). Second, the loss of *GOB* function (*gob-3* mutant) produces underdeveloped carpels (Berger et al., 2009; Fig. S9B). Third, the semi-dominant *GOB* mutant (*Gob-4d*) exhibited gynoecia with ectopic, partially fused carpels, resulting in fruits with extra, malformed locules (Fig. S9A, C, D).

We have previously shown *GOB* transcripts accumulate in 156OE developing ovaries (Silva et al., 2014), linking *GOB* to the miR156/*SlSBP* module. However, we did not find *GOB* transcripts in our transcriptomic data from 1-2-dpi floral primordia, which suggests that *GOB* activation may only occur at later stages of development. Indeed, *GOB* was up-regulated in 6-8-dpi floral buds (when the carpel arises; Xiao et al., 2009) in both 156OE and *pro*, whilst it was down-regulated in 6-8-dpi floral buds from *rSBP15* plants (Fig. S9E). These observations suggested that *GOB* is a common target of *DELLA/PROCERA* and the miR156-targeted *SlSBP15* during the onset of carpel development. Analysis of meristem size in *Gob-4d*/+ floral buds showed no significant differences from WT (Fig. S9F, G), further indicating that the miR156-*SlSBP15-*GA-*GOB* circuity is not associated with FM size, but rather it affects fruit patterning at later developmental stages.

## DISCUSSION

Some aspects of the complex interaction between the age-dependent miR156/*SPL/SBP* module and GA have been relatively well characterized in *Arabidopsis* and tomato, but how GA affects floral meristem activity and fruit patterning in tomato has been unclear. We show that the miR156-mediated regulation of *SlSBPs* is important for controlling floral meristem size, fruit locule number and determinacy. We also found that the fruit development roles of miR156-targeted *SlSBPs*, GA signalling and of the classic *WUS/CLV* pathway are different, but partially overlap. In particular, the loss of *SlSBP* transcripts in 156OE plants had synergistic effects on fruit determinacy when combined with either high GA responses or with reduced activity of *FAS* (the tomato homologue of *CLV3*; Table 1). Synergy generally arises when pathways that converge at a node or hub are disrupted (Pérez-Pérez et al., 2009). The miR156/*SlSBP* hub provides a notable example, given that *SPLs/SBPs* have been shown to connect many unrelated pathways (Wang and Wang, 2015).

Although the genetic interactions between the miR156/*SlSBP* module and GA differed in tomato in terms of floral transition (Silva et al., 2019), their effect on meristem determinacy seems to be broadly conserved in other species. Meristem size is another conserved developmental aspect influenced by both GA and the miR156-targeted *SPLs/SlSBPs*. Tomato *procera/della* exhibited larger shoot and floral meristems, like plants overexpressing miR156 (Fig. 3, S7). The FM phenotype is largely explained by an increase in cell size and cell number in the L1 layer. Because we analysed only the morphology of cells in the L1 of FMs, variations observed in this cell layer may originate from additional morphological adjustments in the inner cell layers of the meristem. Although we did not analyse this conjecture in more detail, de-repression of the miR172-targeted *AP2* in the central zone (CZ) and organizing center (OC) substantially increased *Arabidopsis* inflorescence meristem size (Sang et al., 2022). MiR172 transcripts levels were reduced in the enlarged 156OE floral primordia, whereas miR172-targeted *SlTOE-like* was up-regulated. On the other hand, *SlTOE-like* was similarly expressed in *pro* and WT primordia (Fig. 4, 7), suggesting that miR156-targeted *SlSBPs* regulate the miR172/*TOE-like* node to modulate tomato FM size independently of GA.

On the other hand, overexpressing *rSBP15* in tomato attenuated GA responses in the vegetative development and in the establishment of locule number (Fig. 6, S6). Moreover, high levels of *rSBP15* reduced the number of indeterminate fruits in the *pro* mutant (Fig. 6), indicating that the miR156/*SlSBP15* node and GA interact in distinct developmental contexts in tomato. Similar observations were reported for rice, in which the overexpression of *OsSPL14* blocks GA effects on seed germination, seedling growth and disease susceptibility (Liu et al., 2019; Miao et al., 2019). Loss of miR156 function in *Arabidopsis* leads to the production of a smaller vegetative shoot apical meristem (SAM), whereas the quintuple mutant *spl2;9;10;13*;*15*, which disrupts miRNA156-targeted *SPL* genes, exhibits a larger SAM. In *Arabidopsis*, the miR156-targeted *SPLs* seem to control SAM size by promoting *WUS* expression independently of *CLV3* signalling (Fouracre and Poethig, 2019). However, our molecular and genetic data suggest that in the FM, *SlSBPs*, GA and the *SlWUS-SlCLV3* module converge on shared downstream targets. Future studies on downstream targets such as *SlCRCa* and *SlMBP18* may reveal how the different pathways are integrated at the molecular level, both in the SAM and FM.

In summary, our observations indicate that the miR156/*SlSBP* module functions in tomato fruit development, but its interaction with GA signalling differs between vegetative and reproductive development. Furthermore, the miR156/*SlSBP* module interacts with both GA signalling and with the *WUS/CLV* pathway in ways that differ between developmental stages, from the regulation of FM size to setting locule numbers and the regulation of fruit determinacy. Thus, the interactions between these regulatory modules cannot be easily extrapolated between developmental contexts. Importantly, our work suggests that multiple genetic pathways are available to modify features of tomato fruit development that have so far been associated with the traditional *CLV-WUS* circuity. For instance, although *GOB* is another molecular link connecting gibberellin and the miR156/*SlSBP* module, the higher number of locules observed in the *Gob-4d* fruits is not a result of larger FMs, but rather modifications in the establishment of boundaries during carpel patterning (Fig. S9). Further studies on molecular mechanisms triggering floral determinacy and further gynoecium patterning will provide valuable information for tomato yield improvement, as the meristem size and final number of locules in the mature fruit are key factors for the establishment of the fruit size and shape. The accumulating knowledge of meristem regulatory pathways, and their relevance in regulating crop yield, might contribute to food security and sustainable agriculture in the next decades.

## Material and Methods

### Plant material and growth conditions

All genotypes of tomato (*Solanum lycopersicum*) described in this work were in the cultivar Micro-Tom (MT) background, which was used as wild-type (WT). The *procera* mutant (*pro*), plants MT overexpressing the miR156 (156OE), the miR156-resistant *SlSBP15* allele (*rSBP15*), the *p35S::GA20ox* construct (GA20oxOE), and *sbp15^CRISPR^* plants were previously described (Garcia-Hurtado et al., 2012; Silva et al., 2014; Silva et al., 2017; Silva et al., 2019; Barrera-Rojas et al., 2022). The *fasciated* (*fas*), *GOBLET 4-d* (*Gob4-d*), and *gob-3* alleles were introgressed into MT as described by Carvalho et al. (2011). Plants were grown as described by Silva et al. (2019). Floral primordia at 1-2 days post-inflorescence (dpi) and closed flower buds at 6-8 dpi were collected.

### Crossings

For crosses, flowers were emasculated and manually pollinated 2 days before anthesis to prevent self-pollination. Most crosses were evaluated in the F1 generation, except for 156OE;*pro*, r*SBP15*;*pro*, 156OE;*fas*, r*SBP15*;*fas*, and *pro;fas*. These double mutants were evaluated in the F2 generation, in homozygous plants for the recessive allele *pro* and *fas*.

### Fruit measurements

Fruits were inspected for the presence of at least one ectopic fruit-like structure growing from their stylar end, and scored as indeterminate fruits. Normal fruits (without the presence of ectopic or malformed structures) were cut in transverse sections to evaluate the number of locules. Each genotype was characterized by the percentage of the fruits that produced a specific number of locules. Over a hundred fruits were evaluated per genotype.

### *rSBP3* vector construct and plant transformation

Total RNA was extracted from tomato leaves with Trizol reagent (ThermoFisher Scientific), treated with Turbo DNAse (ThermoFisher Scientific) and cDNA was synthesized using ImpromII Reverse Transcriptase (Promega). *SlSBP3* (Solyc10g009080) ORF was amplified and cloned into pENTR D-TOPO (ThermoFisher Scientific). The miR156 recognition site-containing 3’-UTR was removed from *SlSBP3*, therefore generating the miR156-resistant *SlSBP3* allele (*rSBP3*). After sequencing, the cloned fragment was recombined into pk7WG2.0 (Gateway System) in front of the CaMV35S promoter, using the LR Clonase (ThermoFisher Scientific), generating the *p35S::rSBP3* construct. Tomato MT plants were transformed with the *p35S::rSBP3* construct as described (Silva et al., 2014), generating the *rSBP3* plants. At least five transgenic events were obtained, and three were further analysed.

### RNA extraction, cDNA synthesis and qRT-PCR analysis

Total RNA was treated with DNAse and reverse-transcribed to generate first-strand cDNA, as described above. PCR reactions were performed using GoTaq qPCR Master Mix (Promega) and analyzed in a StepOnePlus real-time PCR system (Applied Biosystems). Tomato *TUBULIN* (Solyc04g081490) was used as internal control. Three technical replicates were analysed for three biological samples (each comprising at least 10 floral primordia or closed buds), together with template-free reactions as negative controls. For miRNA quantification, cDNA synthesis and qPCR reaction were performed as described (Varkonyi-Gasic et al. 2007). The threshold cycle (CT) was determined and fold-changes were calculated using the equation 2^-ΔΔct^ (Livak & Schmittgen, 2001). Oligonucleotide sequences are listed in Table S2.

### Differentially Expressed Genes (DEGs) analysis

Floral primordia at 1-2 days post-inflorescence (dpi) were collected from WT and 156OE plants, and immediately frozen in liquid nitrogen. Two total RNA replicates from each genotype were sent to construct RNA sequencing libraries (RNA-seq) and high throughput sequencing (Illumina NovaSeq platform) at Fasteris Co. Ltd (https://www.fasteris.com/en-us; Switzerland). Raw sequencing reads were first cleaned by removing adaptor sequences and low-quality reads with Trimmomatic (Bolger et al., 2014) and BBDuk (Bushnell, 2021), and library duplication was assessed with dupRadar (Sayols et al., 2016). The resulting high-quality reads were mapped to the tomato reference genome (Tomato Genome Consortium, 2012) (ITAG4.0) and transcripts were aligned, assembled, and quantified using the Hisat2 and Salmon packages with default parameters (Kim et al., 2017; Patro et al., 2017). Differential expression analyses between WT and miR156-overexpressing plants were performed with the edgeR package (Robinson, McCarthy and Smyth, 2010). DEGs in 156-OE compared with WT (MT), with adjusted *P* ≤ 0.05 and absolute fold-change ≥ 2.0, were filtered for further analysis.

DEGs were annotated based on the SOLGENOMICS database (version SL4.0 and Annotation ITAG4.0) (Table S1). Enriched GO terms for the DEG list were identified using the goseq package (Young et al., 2010) and the Gene Ontology Consortium database (Gene Ontology Consortium, 2021). The SOLGENOMICS database was used as a reference for the GO analysis. Fisher’s exact test with FDR correction with a cutoff of 0.05 was applied to determine enriched terms (Benjamini and Hochberg, 1995). Raw sequence data from this study have been deposited in Gene Expression Omnibus (GEO) of NCBI under the accession number GSE223674.

### *In Situ* hybridization

*In Situ* hybridization was performed following the protocol described by Javelle et al. (2011). Primordia from 10-days post-germination (dpg) WT seedlings and pre-anthesis WT ovaries were collected and fixed in 4% (w/v) paraformaldehyde. After alcoholic dehydration, plant material was infiltrated and embedded in Paraplast X-Tra (McCormick Scientific). *SlSBP15* (Solyc10g078700) probe was generated by linearizing (using the *BspHI*; NEB) the pGEM vector containing a 597-bp *SlSBP15* fragment (nucleotides 523 to 1119 of the coding sequence). *In vitro* transcription was performed using the DIG RNA Labeling Kit (SP6/T7, Roche). The sense *SlSBP15* probe was used as a negative control. Locked nucleic acid (LNA) probes with sequence complementary to miR156 (5′-GTGCTCACTCTCTTCTGTCA-3′) and a negative control (scrambled miR, 50-GTGTAACACGTCTATACGCCCA-30) were synthesized by Exiqon (http://www.exiqon.com/), and digoxigenin-labeled using a DIG oligonucleotide 3′ end labeling kit (Roche Applied Science). Ten picomoles of each probe were used for each slide. All hybridization and washing steps were performed at 55°C as described by Javelle et al. (2011). Pictures were photographed using light Microscope Axio Imager.A2 (Carl Zeiss AG). Oligonucleotide sequences are listed in Table S2.

### Hormone treatments

Gibberellin (GA_3_; Sigma-Aldrich; 10^−5^ M), paclobutrazol (PAC) (Sigma-Aldrich; 10^−6^ M), and mock solutions were applied to plants by watering as described (Silva et al., 2019).

### Confocal imaging and image analysis

At least five reproductive floral primordia at 1-2 days dpi were dissected for each genotype, and the first floral meristem was selected in stereomicroscope for a standardized stage. Primordia bearing early floral meristems with sepal primordia were selected, stained by modified Pseudo-Schiff with Propidium Iodide, and imaged as previously described (Bencivenga et al., 2016; Serrano-Mislata et al., 2017).

For image analysis, Python scripts and Fiji macros were used to segment confocal image stacks, define the position of cells within the floral meristem, and to delimit meristem cells and measure L1 volume. The function of each script and instructions on how to perform analysis with them were described in detail in Bencivenga et al. (2016) and Serrano-Mislata et al. (2017).

For SAM area measurements, shoot apices were collected at 4-6 days post-germination and photographed with a Leica stereo microscope S8AP0 (Wetzlar, Germany), coupled to a Leica DFC295 camera (Wetzlar, Germany). Quantification of the SAM area was done in the ImageJ software (NIH). We measured meristem width by drawing a horizontal line between the insertion points of the two youngest visible leaf primordia. From this line, we drew a vertical line to the top of the dome of the meristem to estimate its length (Xu et al., 2015; Rodriguez-Leal et al., 2019). At least four meristems were used for area measurements.

## Statistical analysis

The data are presented as bars and box-plots obtained with GraphPad Prism (https://www.graphpad.com). All statistical analysis was performed using GraphPad Prism. Student’s t test for unpaired data was used. For meristem analysis, ANOVA followed by Tukey’s pairwise multiple comparisons was performed. P values greater than 0.05 were reported as not significant.

## Acknowledgments

L.F.F. received a fellowship from Coordination for the Improvement of Higher Education Personnel (CAPES), Brazil. This work was supported by FAPESP (grant no. 18/17441–3) and Biotechnology and Biological Sciences Research Council (BBRSC no. BB/P013511). M.H.V. and J.P.O.C. were recipients of The São Paulo Research Foundation (FAPESP) fellowships nos. 19/20157-8 and 18/13316-0, respectively. We thank the members of Dr Nogueira’s laboratory for helpful discussions.

## Author contributions

L.F.F., M.H.V., J.P.O.C. and F.T.S.N. were responsible for the conception, planning and organization of the experimental time line. L.F.F., M.H.V., J.P.O.C., C.H.B-R., E.M.S., G.F.F.S., and A. C. Jr carried out experiments. F.T.S.N. and R.S. directly supervised the development of the experimental plan. G.B.A. and G.R.A.M. helped to analyse the RNA-seq data. L.E.P.P. provided some genetic materials. L.F.F., M.H.V., J.P.O.C., R.S. and F.T.S.N. discussed the resulting data. The manuscript was written by L.F.F., R.S. and F.T.S.N.

## Data Availability

The RNA-Seq data underlying this article are available in Gene Expression Omnibus (GEO) of NCBI under the accession number GSE223674. All primary data to support the findings of this study are openly available upon request.

## Summary statement

We show here that tomato floral meristem activity and fruit development are both orchestrated by the interplay between gibberellin and the miR156-targeted *SlSBPs*.

## SUPPLEMENTARY INFORMATION

**Fig. S1.** miR156-overexpressing plants (156OE) showed higher fruit indeterminacy when treated with gibberellic acid (GA_3_).

**Fig. S2.** Examples of 3-D views of raw confocal images of floral meristems stained by the modified pseudo-Schiff propidium iodide (mpS-PI) method.

**Fig. S3.** Expression pattern of miR156 in tomato vegetative and reproductive developmental phases.

**Fig. S4.** Molecular and phenotypic characterization of *rSBP3* transgenic lines.

**Fig. S5.** High expression of a miR156-resistant *SlSBP15* allele reduces meristem volume.

**Fig. S6.** Expression of a miR156-resistant version of *SlSBP15* (*rSBP15*) attenuates gibberellin responses in tomato.

**Fig. S7.** Tomato shoot apical meristem (SAM) activity is modulated by the miR156/*SlSBP15* node, *DELLA/PROCERA* (*PRO*) and *FASCIATED* (*FAS*).

**Fig. S8.** Tomato miR156/*SlSBP* module and *DELLA/PROCERA* act synergistically with *FASCIATED* (*FAS*) to regulate fruit patterning.

**Fig. S9.** *GOBLET* (*GOB*) mis-expression leads to later defects in fruit patterning.

**Table S1.** Differentially expressed genes (DEGs) identified in miR156-overexpressing plants (156OE) floral apices compared with WT.

**Table S2.** Oligonucleotide sequences used in this work.

